# SERS Analysis Of Sialic Acid In Single Dendritic Cells Within The Tumor Microenvironment

**DOI:** 10.1101/2024.08.22.608950

**Authors:** Xingrui Li, Lingling Ma, Xiaofei Yang, Nian Wang, Shiyu Bai, Qiongzhen Zhao, Yu Yang, Weidong Huang, Zhengding Su, Jinyao Li

## Abstract

The crucial role of Dendritic Cells (DCs) in anti-tumor immune responses depends on surface Sialic Acid (SA). The regulation of DC surface sialic acid in the Tumor Microenvironment (TME) remains underexplored. Current methods struggle to provide highly sensitive, multiplex analyses of SA and other immune protein changes in single DCs within the tumor microenvironment. Here, we employed a SERS tags method for specific and highly sensitive analysis of SA, MHCII, CD86, and CD40 in DCs using a single DC analysis microfluidic platform. We also explored the differential regulatory effects of various immune-modulating drugs and tumor supernatant on multiple immunophenotypes of DCs in different immune states. The expanded two-cell microfluidic system further allows for phenotypic analysis of DCs at different time points within the tumor microenvironment. Given this method enables highly sensitive single-cell analysis of DCs, further development of this technology for tumor microenvironment applications will aid in deeply understanding the tumor-induced suppression of DC immune function, providing valuable insights for DC-mediated tumor immunotherapy research.

## 1. Introduction

Sialic Acid (SA) is typically the terminal monosaccharide of glycans on cell surface glycoproteins^1^. SA mediates various immune processes, including host-pathogen recognition, cell migration, and antigen presentation. SA mediates various immune processes, including host-pathogen recognition, cell migration, and antigen presentation. SA plays a significant role in immune responses, including cellular immune recognition. It is a key determinant of cell surface receptors and regulates the function and differentiation of immune cells through protein glycosylation^2^. Consequently, precise analysis of SA expression changes on the surface of immune cells is essential for advancing immune cell research^3^.

Dendritic cells (DCs) are key antigen-presenting cells that deliver ingested antigens to T lymphocytes and regulate both cellular and humoral immune responses^4^. Upon antigenic stimulation, DCs differentiate into mature cells, exhibiting upregulated expression of co-stimulatory molecules, including surface MHC II, CD86, and CD40^5^. Within DC antigen-presenting functions, SA is believed to regulate critical factors essential for this role. Removal of surface sialic acid from DCs using sialidase significantly enhanced bacterial phagocytosis, promoting functional maturation and increasing immunoreactivity of DCs^6,7^. Another critical immune function of DCs is chemotaxis toward lymph nodes for antigen presentation. CCR7, the primary chemokine receptor driving DC migration, facilitates the delivery of antigens to lymph nodes. After sialic acid removal from CCR7, DCs lost chemotactic ability, primarily because sialic acid serves as a binding site regulating CCR7-CCL21 interaction^8,9^. Without affecting DC differentiation and survival, a sialic acid synthesis inhibitor enhanced T cell proliferation and affinity interactions between DCs and T cells, particularly in antigen-dependent interactions^10^. Without affecting DC differentiation and survival, a sialic acid synthesis inhibitor enhanced T cell proliferation and affinity interactions between DCs and T cells, particularly in antigen-dependent interactions^11-13^. This insight offers a pathway to enhance cellular immunotherapy, particularly in tumor immunotherapy.

Various approaches have been developed to detect SA, including colorimetry, electrochemistry, mass spectrometry, and fluorescence^14-17^. Quantitative detection of SA expression on intact cell surfaces remains a challenge. These methods for analyzing SA on living cells still face issues such as poor accessibility, activity disruption, and reliance on additional signal amplification techniques. Surface-enhanced Raman scattering (SERS) is widely utilized in bioanalysis for its non-destructive and highly sensitive properties^18-20^. Due to cellular complexity, non-targeted SERS methods struggle to obtain SA results without interference^21^. Thus, there is an urgent need to develop SA-targeted SERS tags to avoid biological system interference and obtain pure SA data^22^.

This study combines microfluidics with SERS-tag technology, offering a novel approach to understanding the relationship between SA changes and immune function in DC cells. We propose a microfluidics-based strategy for SERS analysis of SA on individual live DCs. The SERS-tag employs MPBA-labeled AuNPs, demonstrating high selectivity for SA under physiological conditions. MBN serves as a targeting molecule with a Raman reporter, enabling the SERS-tag to bind to DC surfaces and determine SA expression levels based on MPBA signals. Additional SERS-tags targeting DC functional proteins (MHCII, CD40, CD86) were prepared, allowing simultaneous analysis of DC immunophenotypes and SA, revealing a negative correlation between SA and immune molecule expression in DCs. Using this SERS-tag, we investigated a microfluidic system to observe SA up-regulation in response to tumor influence within a complex single-DC/single-tumor environment. Our results demonstrate the capability of highly sensitive SA analysis on individual DCs within the SERS-based microfluidic system and reveal differences in SA expression in DCs influenced by tumors. This method provides a promising approach for highly sensitive analysis of SA changes in immune cells within the tumor microenvironment.

**Scheme 1.**
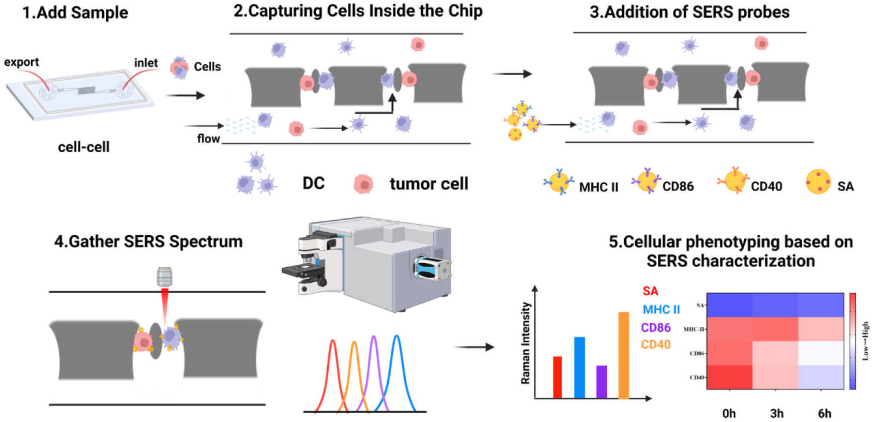
Workflow of the microfluidic device-SERS approach for multi-pathway analysis of the phenotype of single-cell DCs.

## 2. Experimental section

### 2.1 Materials and reagents

Hydrogen tetrachloroaurate trihydrate (HAuCl_4_ 3H_2_O), N-hydroxysuccinimide (NHS), 4-Mercaptophenylboronic acid (MPBA) were purchased from Tianjin Hiens Biochemical Technology Co. (Tianjin, China). N-(3-dimethylaminopropyl)-N ′-ethyl carbodiimide hydrochloride (EDC), 5,5’-dithiobis (2-nitrobenzoic acid) (DTNB), 7-mercapto-4-methylcumarin (MMC) were purchased from Shanghai Hohong Biomedical Technology Co. (Shanghai, China). 4-mercaptobenzoic acid(4-MBA) was purchased from Shanghai McLean Biochemical Technology Co. (Shanghai, China). Trisodium citrate, anhydrous ethanol (CH_3_CH_2_OH, analytical purity) were purchased from Xinbute Chemical Co. (Tianjin, China). Fetal bovine serum (FBS) was purchased from Biological Industries, Inc. (Palestine). Trypsin was purchased from All Style Gold Biotechnology Co. (Beijing, China). RPMI Medium 1640, phosphate buffer saline (PBS) 、 penicillin-streptomycin (PS) were purchased from Thermo Fisher Scientific (Waltham, MA, USA). Granulocyte-macrophage colony stimulating factor (GM-CSF) was purchased from Peprotech (USA). Anti-MHC II , anti-CD86,and anti-CD40 were purchased from Elabscience (Wuhan, China). Lipopolysaccharide (LPS), α2-3,6-Neuraminidase, Clostridium perfringens, Recombinant, E. coli were purchased from Sigma-Aldrich (St. Louis, MO, USA). SU-8 3050 Photoresist, PDMS prepolymer, SU-8 photolithographic developer, silicon chip were purchased from Semiconductor Manufacturing International Corporation. (Suzhou, China). In all experiments, ultrapure deionized (DI) water (Milli-Q water purification system, 18.2 MΩ) (Millipore Corp., Burlington, MA, USA) was used.

### 2.2 Instrumentation

UV-visible absorption spectra were recorded using a NanoDrop™ One Ultra-Micro UV Spectrophotometer (Thermofisher, USA), and morphological characterization was performed using a JEM-2100 transmission electron microscope (TEM, JEOL, Tokyo, Japan). The particle size and zeta potential of AuNPs and nanoprobes were measured using a Zetasizer Lab ZS90 nanoparticle sizer analyzer (Malvern, UK). Confocal microscopy images were taken using Nikon AIR+Ti2-E, DC cell biomarker expression was analyzed using a CytoFLEX flow cytometer (Beckman, Germany), a laser confocal microscope (Nikon, Japan), and SERS measurements were carried out on a Horiba LabRAM Odyssey high-speed, high-resolution microscopic confocal Raman spectrometer in the The system was calibrated at 520.7 cm-1 using in silico prior to testing.

### 2.3 Preparation of gold nanoparticles

The sodium citrate reduction method was used to prepare the nanogold solution^23^. Add 1 mL of 1% concentration of chloroauric acid solution into a round bottom flask to 100 mL, place in a constant temperature heating magnetic stirrer heated to boiling, and then 0.5 mL of 1% sodium citrate solution was added quickly, the visible color will be transparent to gray-black, and then finally become a clarified and transparent violet-red. The reaction continued to heat and stir for 20 min until the reaction solution no longer changed color, stopped heating and cooled to room temperature, put into the refrigerator at 4°C to keep away from light.

### 2.4 Preparation of specific immune SERS-tag

Specific immuno-SERS labeling by surface modification of AuNPs. To prevent AuNPs from polymerizing during surface modification, 10 μL of Tween 20 was first added to 2 mL of AuNPs solution and stirred rapidly at room temperature for 30 min. 2.0 mL of AuNPs solution was then mixed with 4-MBA, DTNB, and MMC Raman molecules (10 μL, 1 mM ethanol solution) and stirred for 2.0 h. The Raman molecules were attached to the AuNPs, and then centrifuged at 6000 rpm for 10 min to remove unbound free Raman molecules, which were then dispersed into PBS, and 5.0 μL of EDC (1 mM) and NHS (1 mM) were added respectively and left to react for 1 h. Next, 2 μg of MHC-II, CD86, and CD40 antibodies were mixed into a mixture. II, CD86, and CD40 antibodies were added to 4-MBA, DTNB, and MMC-labeled AuNPs, respectively, and incubated at 4°C overnight. Finally, the antibody-coupled nanoprobes were centrifuged at 6000 rpm for 15 min to remove the unreacted free antibody, and the centrifuged substrate was re-dispersed in 0.1% (W/V) BSA to obtain the specific immuno-SERS nanoprobes, which were stored in the refrigerator at 4°C.

### 2.5 Preparation of salivary acid SERS nanoprobes

2.0 mL of AuNPs solution was mixed with Raman molecules MPBA (10 μL, 1 mM ethanol solution), and stirred for 2.0 h. The Raman molecules were modified onto the AuNPs, and then the unbound free Raman molecules in the mixture were removed by centrifugation for 10 min at 6,000 rpm, the supernatant was removed, and then dispersed into PBS and placed in the refrigerator at 4°C for backup. SERS nanoprobes were obtained to recognize salivary acid.

### 2.6 DC induction and culture of bone marrow-derived

Tibia and femur were removed from C57BL/6, and the bone marrow was flushed with RPMI 1640 medium, and the bone marrow cells were collected and cultured in RPMI-1640 medium containing 10% inactivated fetal bovine serum, 1% streptomycin-penicillin, and 20 ng/mL GM-CSF, and were placed in a 37°C cell culture incubator with a CO2 concentration of 5%. The fluid was half changed on day 2, full changed on day 3, half changed on day 5, and the cells were collected on day 6, which were the immature DCs from the induced culture.

### 2.7 Tumor cell culture

CT26 colon cancer cells were cultured in RPMI-1640 medium (containing 10% fetal bovine serum, and 1% streptomycin-penicillin) and placed in a 37°C cell culture incubator with a CO2 concentration of 5%. Cells were collected under the cell density grew to 80%-90%.

### 2.8 Preparation of tumor cell culture supernatant (TCM)

Routinely cultured colon cancer CT26 cells, collected CT26 cells in the logarithmic growth phase, inoculated in complete medium at a concentration of 1×10^4^ cells/mL, placed in 37 °C, 5% CO2 incubator, collected cell culture supernatant of passaged culture for 48h, filtered by microporous filter membrane (0.22 μm), and frozen at -80 ° C for spare.

### 2.9 Acquisition of SERS spectral data

SERS spectra were obtained on a confocal Raman system (LabRAM Aramis, Horiba Jobin Yvon). A laser (wavelength 633 nm) was used in the detection with a laser power of about 25%. The samples were calibrated using a silicon wafer at 520.7 cm ^-1^ before detection. For all cell samples in this experiment, the integration time of the SERS assay was 5 s, accumulated twice.

## 3. Results and Discussions

### 3.1 Design and characterisation of SERS-tag nanoprobes

To analyze SA on single dendritic cells and multiple immunophenotypes, various SERS-tags were designed and characterized. The synthesis process of SERS nanoprobes is illustrated in Fig. 1A. AuNPs with an average diameter of approximately 68 nm were prepared (Fig. 1B). 4-MBA, DTNB, and MMC were selected as Raman reporter moieties to modify AuNPs with antibodies of different specificities (MHC II, CD86, and CD40). The zeta potential changes of AuNPs during modification with signaling molecules and antibodies were determined to be -42, - 20.8, and -9.2 mV, respectivel, and the plasmon absorption peaks exhibited a red shift. The characteristic SERS signals of the prepared AuNPs@4-MBA were strong, and no significant difference was observed in the SERS signals of AuNPs@4-MBA@Ab before and after antibody modification, indicating that the antibody modification process did not alter the SERS signals (Figure 2C). These results confirm the successful preparation of the SERS-tag. MPBA, which possesses both sialic acid targeting and Raman reporter functions, was selected to modify AuNPs for targeting sialic acid^24^. For SERS analysis of multiple dendritic cell phenotypes, the Raman characteristic peaks of the four SERS-tags were selected as 1423 cm^−1^ (4-MBA), 1334 cm^−1^ (DTNB), 1175 cm^−1^ (MMC), and 998 cm^−1^ (MPBA), respectively, to avoid peak overlap (Figure 1D). The signals from 10 batches of SERS-tags were measured, yielding a coefficient of variation (CV) of 8.8%, demonstrating excellent reproducibility. The prepared multi-target probes were subsequently used for single-cell biomarker analysis.

**Fig 1.**
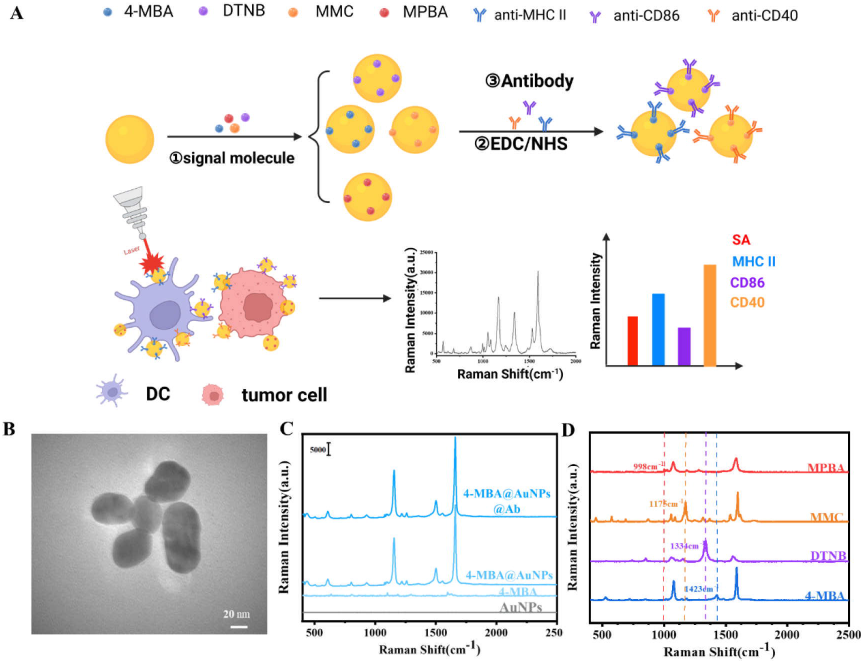
Design of SERS-tag with multi-pathway analysis principle (A) Design of different SERS-tag nanoprobes. (B) TEM image showing the morphology of AuNPs as a 68 nm diameter circle. (C) SERS spectra of AuNPs, 4-MBA, 4-MBA@AuNPs, and 4-MBA@AuNPs@Ab. (D) The four SERS-tag nanotag feature peaks are independent of each other.

**Fig 2.**
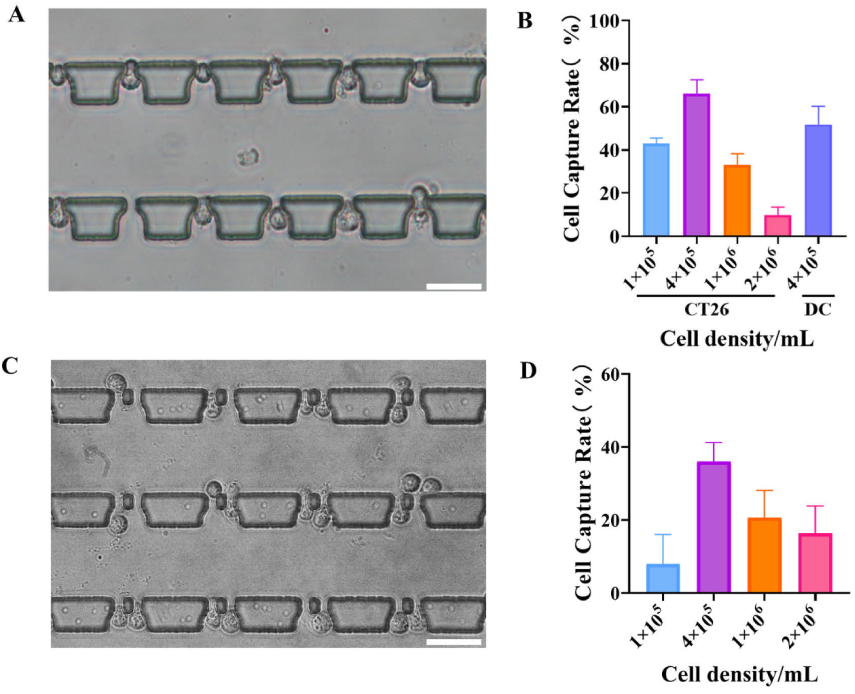
(A) Single cell capture of dendritic cells (B) Single cell capture efficiency at different cell concentrations. (C) Two cells captured within microfluidic structure. (D) Efficiency of two-cell enviroment with different cell concentrations.

### 3.2 Acquisition of individual dendritic cells in microfluidic devices with multiple immunophenotyping analyses

In order to achieve multiple phenotyping analyses of single cell DCs within microfluidic chips, a simple and efficient single cell capture system was designed using microfluidics. First, the cell concentration was adjusted to optimize the throughput of single cell and two-cell from the suspension. Capture efficiency was optimized using CT26 cells in the chip. At a flow rate of 50 μL/h, different cell concentrations (1×10^5, 4×10^5, 1×10^6, and 2×10^6 cells/mL) were injected into the chip, yielding capture efficiencies of 41.3±3.5%, 66.7±7.5%, 32.7±4.5%, and 10.4±6.4%, respectively. Under the same conditions, the capture efficiency for dendritic cells was 51.5±5.5%, demonstrating that the system could effectively isolate single dendritic cells for phenotypic analysis. Under the same conditions, two-cell capture chambers were designed to facilitate rapid acquisition of single DCs and tumor cells within the tumor microenvironment. The efficiency of two-cell structure to capture two cells under identical fluid conditions was 7.8±8.1%, 36.7±5.3%, 21.2±7.3%, and 16.4±6.9%, respectively (Figure 2C-D). At higher cell concentrations, multiple cells were captured in one chamber; at lower concentrations, longer capture times were needed to isolate two cells. The entire capture process was completed within 30 minutes, given the rapid immunoregulation of DCs. A cell concentration of 4×10^5 cells/mL was selected as the optimal condition for efficient two-cell capture using the microfluidic chip.

To analyse the phenotypic heterogeneity of single-cell DCs within the microfluidic chip, we first intrachip the surface molecular detection process for individual DCs as shown in Figure 3A. The flat Raman spectra of the spit detection probes for each group of collected single DCs are shown in Fig. 3B. The different peak shifts of the Raman spectra corresponded to different target molecules, 1423 cm-1 (MHC II), 1334 cm-1 (CD86) and 1175 cm-1 (CD40). We used the classical LPS-stimulated DC maturation method to obtain mDC while using tumour (CT26) medium supernatant (TCM) treated different iDCs and mDCs. compared the Raman spectroscopy results of the cells after different treatments. Compared with untreated iDC, SA of mDC was significantly down-regulated, while other immune molecules were significantly up-regulated. It indicated that with the maturation process of DC, the immune function of DC was enhanced and the expression of sialic acid of DC was decreased.TCM had a significant inhibitory effect on the immune maturation of DC, while the analysis of sialic acid revealed that TCM maintained the sialic acid level of DC at a high level (Figure 3B).

**Fig 3.**
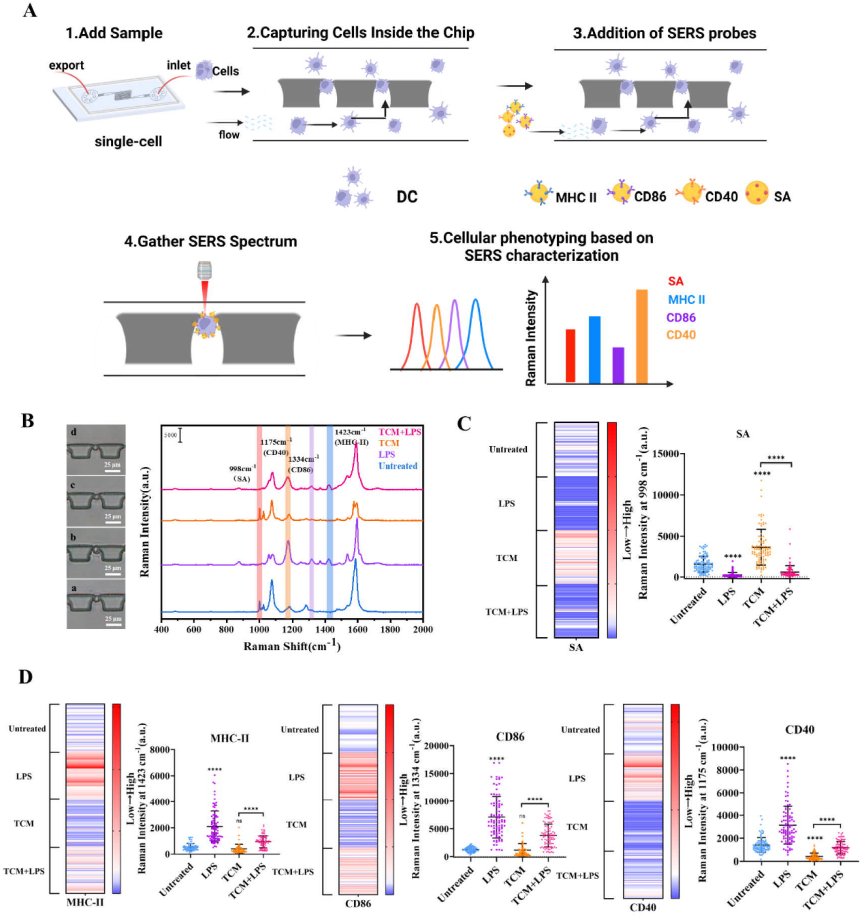
Heterogeneity analysis of SERS Raman spectra of single DCs obtained within the microfluidic device. (A) Workflow of microfluidic device to capture single DC and phenotypic multi-analysis of single DC using SERS-tag. (B) Raman spectra of different treated DCs. The scale bar is 25 μm.(C) The results of 90 single DCs with SA expression profiles analysed using Raman spectroscopy. (D) Raman spectroscopy results of DCs at 1423 cm-1 (MHC II), 1334 cm-1 (CD86) and 1175 cm-1 (CD40).

Further analysis of SA in single DCs involved counting 90 single DCs from each group (Fig. 3C,). Heterogeneity was observed in the surface functional molecules and SA among individual DCs. Compared to mean value analysis, single-cell statistics more effectively demonstrate differences among individual DCs. Single-cell analysis revealed that DCs stimulated with LPS showed significantly down-regulated surface SA. This is due to the fact that DC mature by down-regulating SA which in turn reduces the ability to recognise and phagocytose external antigens in order to gain the ability to deliver antigens to T cells. This was confirmed by analyzing other immunocompetent molecules on the DCs. TCM significantly affected DCs by upregulating sialic acid, thereby inhibiting the maturation process. Despite an increase in immune molecules (MHCII, CD86, CD40) following LPS stimulation, SA expression was downregulated. However, this downregulation was not significantly different from that observed in iDCs. Therefore, the TME can inhibit antigen presentation by upregulating SA in DCs and inhibiting DC maturation (Figure 3D).

To demonstrate the applicability of single-cell SA analysis in assessing DC responses to drugs, two functionally distinct DC-regulating drugs were selected to analyze single DC responses to different treatments (Figure 4A). In a previous study, we found that glycyrrhiza glabra polysaccharide (GUPS) significantly promotes DC immune maturation and activates DCs to drive anti-tumor immunity via the TLR4-NF-κB pathway^25,26^. TPCK, a commonly used NF-κB pathway inhibitor, was employed as a DC immune function suppressor in this study^27^. Within the microfluidic device, the affinity of SERS-tags for DC surface receptors mirrored population-level trends but exhibited significant heterogeneity among single cells. In GUPS-treated DCs, SA expression significantly decreased, while the 3 kinds of immune proteins (MHCII, CD86, and CD40) increased significantly, which was consistent with our previous results. The TPCK inhibitor caused upregulation of SA and downregulation of immune function proteins on DCs. These suggests that downregulation of SA is a key indicator of DC maturation. After single-cell capture by the microfluidic chip, cell-cell communication was blocked, leading cells to undergo different stages of growth and development. This resulted in greater variability among cells following drug treatment (Figure 4B-C). Significant heterogeneity in single DC responses to immunomodulatory drugs was demonstrated.

**Fig 4.**
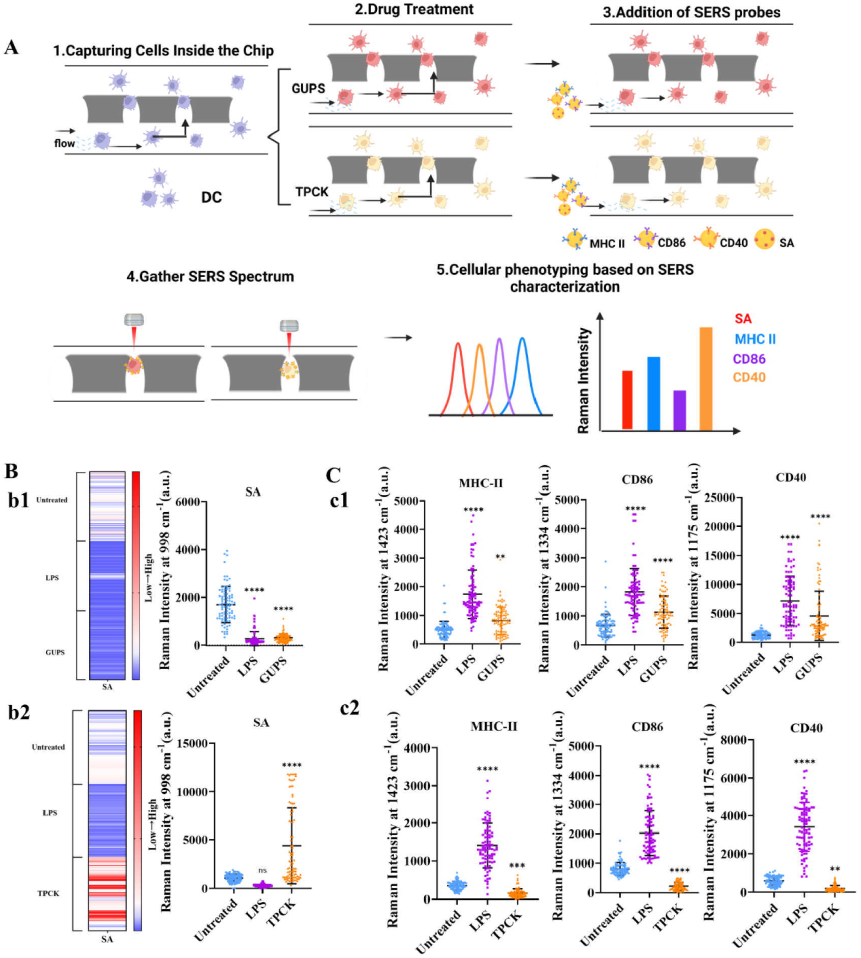
Multiple phenotypic analyses of single DCs after drug treatment. (A) Process of analysis of different drug-treated DCs within the microfluidic device. (B) Raman spectral analysis of single DC after GUPS treatment. (C) Raman spectral analysis of single DC after TPCK treatment. Each set of single cells was analysed using data from 90 cells.

### 3.4 Single cell analysis of tumour microenvironment regulation of DCs

The suppression of DCs by the TME is a critical factor affecting tumor immunotherapy. The use of microfluidic devices to efficiently construct TMEs where tumors and DCs coexist, along with the analysis of individual DC heterogeneity within these TMEs, can help explore the regulatory mechanisms of TME-induced DC immunosuppression. To this end, we designed a microfluidic chamber for dual-cell capture, forming a tumor cell-DC system to study DC immune phenotype and SA changes. Two cell types were introduced into the chip at a flow rate of 50 μL/h at a to the optimised concentration and the ratio (tumor:DC=1:5). The dual-cell chamber captured the two cells separately (Figure 5A). Random combinations of the two cells formed three two-cell systems: DC-DC, tumor-DC, and tumor-tumor. We specifically labeled each cell type with DC-CD40 (APC-red) and CT26-EpCAM aptamer (FITC-green) and observed random combinations of the two cells (Figure 5B). Using the SERS-tag approach, the two cells were analyzed to quickly distinguish between different cell combinations (Figure 5C). This is attributed to the lack of MHCII, CD86, and CD40 expression in tumor cells. This suggests that our constructed two-cell system can generate a single tumor cell-single DC microenvironment, studies on the direct effects of single tumor cells on single DCs.

**Fig 5.**
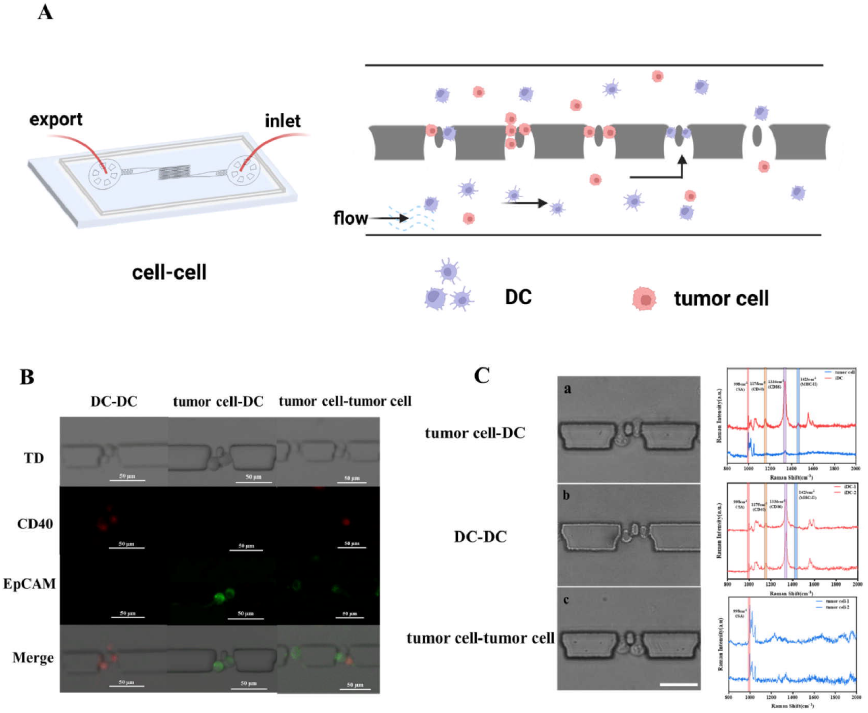
(A) Tumor-DC captured by microfluidic chamber. (B) The three kinds of dual-cell systems. Tumor cells in green and DC cells in red (C) Different species of dual-cell system can be verified for cell species using Raman spectroscopy.Tumor cells in green and DC cells in red (C) Different systems of dual-cell can be verified for cell species using Raman spectroscopy.

To study the biological behavior of individual DCs in the TME and investigate the regulatory effects on DCs, we examined TME-induced regulation of DCs in a time-dependent manner. To study the biological behavior of individual DCs in the TME and investigate the regulatory effects on DCs, we examined TME-induced regulation of DCs in a time-dependent manner. Significant differences in SA expression were observed between iDCs and mDCs co-cultured with tumors across the three time points (Fig 6). iDCs exhibited a significant decrease in SA at 6 h, corresponding to an increase in SA expression in mDCs. We also analyzed different immune protein expressions in DCs. In iDCs, as SA decreased, MHCII expression increased, CD40 expression declined, and CD86 remained unchanged. We also analyzed different immune protein expressions in DCs. In iDCs, as SA decreased, MHCII expression increased, CD40 expression declined, and CD86 remained unchanged. The increase in MHC II on the iDC surface may result from DC immunostimulation due to exocytosis of proteins from tumor cells, which activates antigen phagocytosis, surface antigen processing, and subsequent formation of MHC-antigen peptide complexes. However, the downregulation of CD40 and the lack of changes in CD86 hindered T cell activation, leading to tumor immunosuppression. In mDCs, the tumor suppresses the already upregulated immune function. Based on the experimental results, we speculate that the TME affects antigen phagocytosis in unactivated iDCs but suppresses the function of T cells activated by DCs. For activated mDCs, the TME can exert immunosuppressive effects through various mechanisms, from antigen presentation to T cell activation. Using the two-cell system constructed by microfluidics, Raman spectroscopy visualized the phenotypic modulatory effects of the TME on individual DCs, which plays a crucial role in studying DC-mediated immune responses in TME.

**Fig 6.**
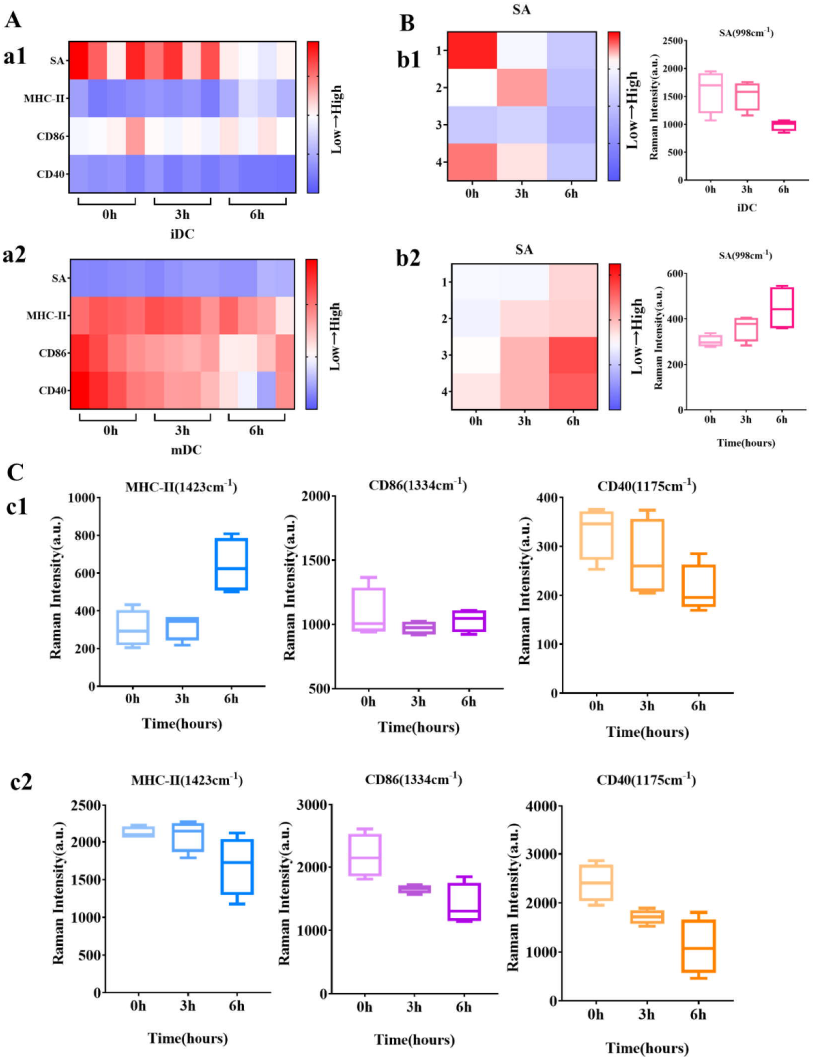
Immunophenotypes of two DCs in a two-cell system with a tumour-DC composition of the TME system analysed by Raman spectroscopy. (A) TME effects on multiple immunophenotypes of iDC and mDC. (B) Effect of TME on SA of iDC and mDC. (C) c1. effect of TME on MHCII, CD86, CD40 of iDC. c2. effect of TME on MHCII, CD86, CD40 of mDC.

## 4 Conclusion

In conclusion, the influence of the TME on individual DCs was studied using a constructed two-cell microfluidic system with the SERS-tag method. The change in SA was identified as key evidence for TME regulation of DC immune activity. AuNPs were used to bind different Raman signaling molecules (4-MBA, DTNB, MMC, MPBA) to form distinct SERS-tags, which were then used to modify MHCII, CD86, and CD40-specific antibodies. A specific SERS-tag for SA was constructed based on the targeting effect of MPBA on SA. Raman spectroscopy enables the rapid analysis of multiple immunophenotypes in single DCs within a microfluidic chip, allowing for the study of regulatory effects of TCM and various immunoregulatory drugs on DCs. Importantly, using the two-cell system, the SERS method enabled time-dependent single-cell level analysis of the immunosuppressive effects on both types of DCs in the TME. Therefore, we anticipate that this method will play a crucial role in achieving more accurate and sensitive analysis of DCs in TME.

## Acknowledgments

Xingrui Li was supported by the Basic Research Projects of Universities in Xinjiang (202406120001), School-Enterprise Cooperation Projects (202399140010, 202306140004). Jinyao Li. was supported by the Tianshan Talent Training Program (2023TSYCLJ0043). Author Contributions: Writing—original draft preparation, X. R. L, N. W. and W. D. H.; writing—review and editing, X. R. L, R. S., Z. D. S. and J. Y. L. All authors have read and agreed to the published version of the manuscript.

## Author contributions

X. R. L., L.L.M., and X.F.Y., designed all experiments and analyzed data. L.L.M, X.F.Y., S.Y.B., Y.Y., and Q.Z.Z., conducted all experiments. X.R.L, L.L.M, N.W.,S.Y.B.,W.D.H., Z.D.S, and J.Y.L, wrote the manuscript.

## Competing interests statement

All authors declare that the research was conducted in the absence of any commercial or financial relationships that could be construed as a potential conflict of interest.

## Notes

### Competing Interest Statement

The authors have declared no competing interest.

